# Targeted RNA NextGenSeq profiling in oncology using single molecule molecular inversion probes

**DOI:** 10.1101/440065

**Authors:** Krissie Lenting, Corina N.A.M. van den Heuvel, Anne van Ewijk, Elizabeth Tindall, Ge Wei, Benno Kusters, Maarten te Dorsthorst, Mark ter Laan, Martijn A. Huynen, William P. Leenders

## Abstract

Hundreds of biology-based precision drugs are available that neutralize aberrant molecular pathways in cancer. Molecular heterogeneity and the lack of reliable companion diagnostic biomarkers for many drugs makes targeted treatment of cancer inaccurate for many individuals, leading to futile overtreatment. To acquire a comprehensive insight in aberrant actionable biological pathways in individual cancers we applied a cost-effective targeted RNA next generation sequencing (NGS) technique. The test allows NGS-based measurement of transcript levels and splice variants of hundreds of genes with established roles in the biological behavior in many cancer types. We here present proof of concept that the technique generates a correct molecular diagnosis and a prognosis for glioma patients. The test not only confirmed known brain cancer-associated molecular aberrations but also identified aberrant expression levels of actionable genes and mutations that are associated with other cancer types. Targeted RNA-NGS is therefore a highly attractive method to guide precision therapy for the individual patient based on pathway analysis.

## Introduction

Many different cancer types share the same driver mutations. Examples are loss of function mutations in tumor suppressor genes (e.g. *p53* and *CDKN2A*, leading to genetic instability and loss of cell cycle control) [1, 2] and DNA repair proteins (leading to genetic instability) [3], and activating mutations, amplifications or fusion events in proto-oncogenes that activate the phosphoinositide 3-kinase (PI3K) and mitogen-activated protein kinase (MAPK) pathways [4, 5]. Other shared features are related to micro-environmental effectors, such as altered metabolism as a result of hypoxia [6, 7] and induction of angiogenesis [8].

Additionally, aberrations exist that are more specific for certain tumor types. Examples are expression of hormone receptors in cancers of prostate, ovary and breast [9, 10]; mutations affecting metabolism (isocitrate dehydrogenase mutations [IDH1 p.R132H] in glioma and acute myeloid leukemia [11]); and mutations affecting the PI3K pathway (KRAS mutations in adenocarcinomas [12], the *BRAF*^*V600E*^ mutation in melanoma [13]). Such specificity is however never absolute. As an example, *IDH-* and *BRAF* mutations are sporadically found in other cancers too [14–16]. Detection of such relatively rare, and therefore *a priori* unexpected mutations in individual patients could lead to repurposing of precision medicines in so called basket trials, in which a precision drug is administered to patients that are recruited based on the identification of an accompanying diagnostic biomarker [17]. Nowadays tumors are increasingly analyzed with cancer hotspot panels to detect aberrations in DNA, either obtained from tumor tissue or from circulating tumor DNA [18–20].

A number of actionable biological pathways in cancer involve the products of genes that are not mutated, but epigenetically regulated, for example by altered transcription factor availability, repressor activity or gene methylation [21, 22]. The activity of such pathways cannot be directly inferred from DNA analyses. Sophisticated machine learning algorithms to analyze whole genome methylation data has been shown to have diagnostic power [23] but this technique is not suitable for reliable pathway analysis. An example is angiogenesis, initiated by hypoxia-induced expression of an abundance of growth factors and followed by extensive crosstalk between tumor cells, tip-and stalk endothelial cells, and pericytes [24, 25]. DNA analysis also does not provide information on post-transcriptional events such as alternative splicing. For example, expression of alternative splice variants of vascular endothelial growth factor (VEGF-A) has important implications for the regulation of angiogenesis [26]. Furthermore, alternative splicing of receptor tyrosine kinases can lead to auto-active and oncogenic PI3K signaling [e.g. EGFR variant III (EGFR^vIII^) and the MET variants MET^Δ7-8^ and MET^Δ14^ [27–30]. To collect this information gene expression data of thousands of cancers are collected in the context of initiatives such as The Cancer Genome Atlas (TCGA). Analysis of these microarray-or next generation sequencing-based datasets has yielded invaluable information on biological pathways in different cancer types but are costwise not suitable for implementation in routine patient care, a necessity to deal with the molecular heterogeneity in cancers. Also, identification of RNA splice variants from RNA next generation sequencing can be difficult because of low coverage of exon-exon boundaries. There is a huge need for methods to obtain such clinically important information for individual tumors in a reliable fashion in order to be able to guide personalized treatment approaches.

Gliomas are primary brain tumors. Due to its low incidence (6/100,000) and high molecular heterogeneity [31] it is a difficult tumor type to organize clinical trials with, although the molecular underpinnings of gliomagenesis and glioma progression are relatively well established. In the absence of alternatives, treatment of its most malignant form, glioblastoma, is still confined to surgery, followed by chemotherapy with temozolomide (TMZ) and radiotherapy [32]. The standard treatment protocols for glioblastoma prolong life expectancy by only a few months. Surgical cure for this tumor type is not possible because complete resection cannot be achieved due to its diffuse infiltrative nature [33]. Glioma is therefore one of the most challenging tumors for which new treatment strategies are urgently needed.

We here present the results from a recently developed technique of quantitative targeted RNA next generation sequencing (t/RNA-NGS), applied on 75 brain tumors, suspected for glioma [34, 35]. The technique which uses single molecule molecular inversion probes (smMIPs) sensitively and quantitatively measures expression levels of potentially actionable genes that play a role in multiple cancer types, and unequivocally identifies splice variants at low costs. We show that with the right choice of target transcripts it is possible to let algorithms calculate an appropriate molecular diagnosis and prognosis. This technique can be applied on individual cancers and may therefore have future value in predicting response to targeted therapies for individual patients.

## Results

### Prognostic value of t/RNA-NGS profiles

To investigate the prognostic value of t/RNA-NGS datasets we profiled a cohort of 75 brain tumors, including 69 grade II-IV diffuse gliomas and 6 brain lesions that upon routine histopathology were diagnosed as an ependymoma (n=1), one dysembryoplastic neuroepithelial tumor (DNET), one ganglioglioma, a variant glioma, a brain metastasis of lung adenocarcinoma and a lymphoproliferative disorder (LPD) (Table SI). Annotated unique smMIP counts for each tumor sample ranged from 275,000 to 1,111,000 (not shown). Hierarchical unsupervised agglomerative clustering of the gene expression data of the grade II-IV gliomas (excluding the 6 rare cancers) resulted in 3 main clusters A, B and C, comprising of 26, 38 and 5 tumors, respectively (Fig. 1).

**Fig. 1:**
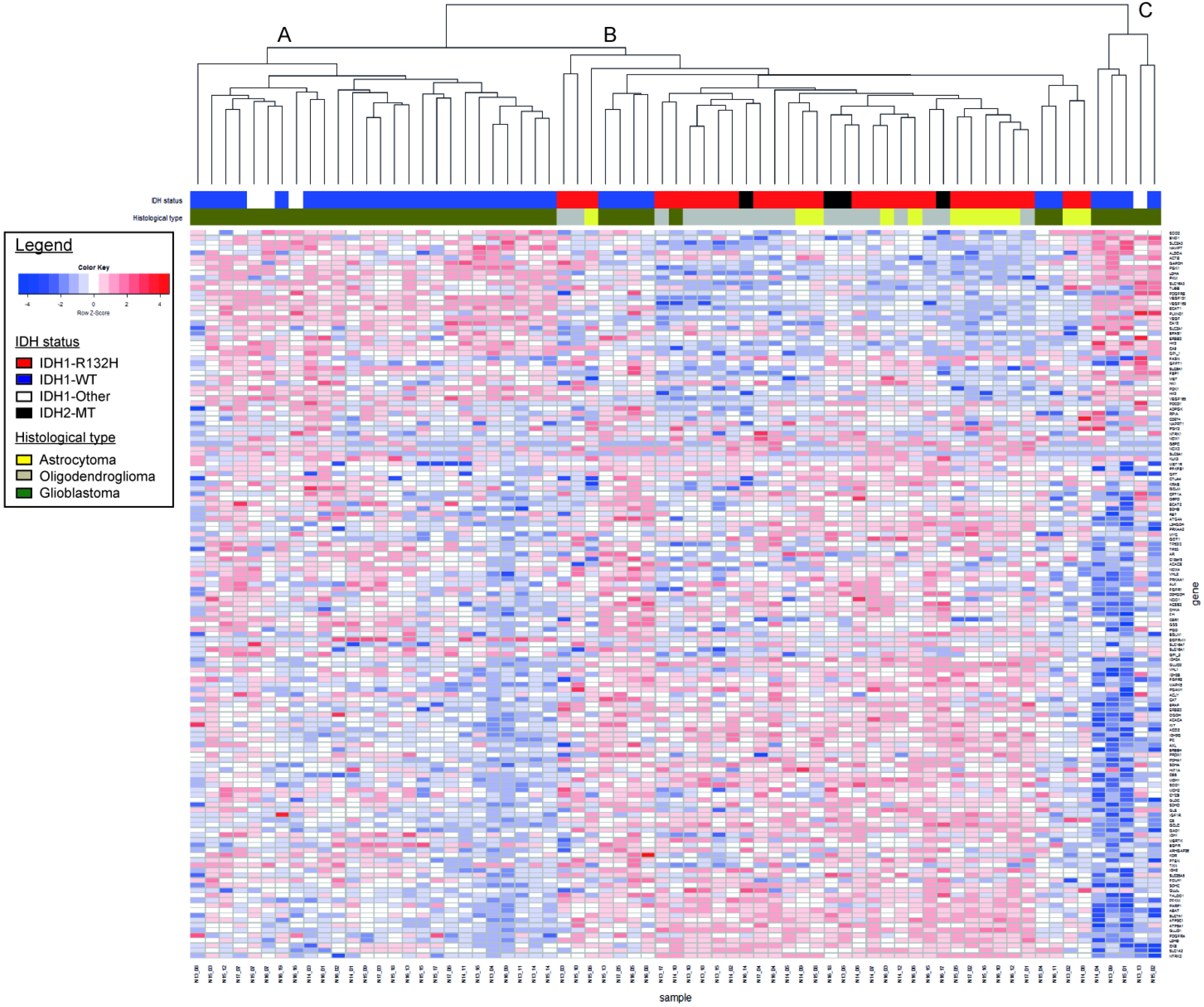
Agglomerative clustering of t/RNA-NGS profiles of 69 grade II-IV gliomas. **C**lustering was based on expression levels of 145 genes of interest. Gene expression levels (in FPM) were transformed to a z-score for each individual transcript. After generating the dendrogram and heatmap using the average clustering method in R, histopathology results and mutation status were added in retrospect (legend and upper annotation bars). Cluster A and C strongly correlate with a wild-type IDH status, while cluster B is strongly associated with the IDH1^R132H^ mutation. IDH2 mutations (R172K, R172M, and R172W) are annotated as IDH2-MT (black in the annotation bar) and all coclustered with the IDH1^R132H^ mutation. IDH1-other refer to variants other than the IDH1^R132H^ (see Table SI).

In a next step we performed a Wilcoxon Mann-Whitney U test to find genes that were differentially expressed between clusters A and B; A and C; and B and C. A total of 83 genes were differentially expressed between clusters A and B with p<0.05 and False Discovery Rate (FDR)<0.05. Among these were transcripts encoding transporters and enzymes involved in metabolism [35]. Membrane receptor tyrosine kinases (RTKs) NTRK2, ERBB3, ERBB4 were higher expressed in cluster B with average fold changes (FC) of 3.5, 4.8 and 5.2 respectively, while cluster A was characterized by significantly higher expression levels of EGFR^vIII^ (FC=250) and VEGF-A isoforms VEGF-A_121_, VEGF-A_165_ and VEGF-A_189_ (all with FC>11, Table SIIa). Between cluster B and C 69 genes were differentially expressed (Table SIIb). Cluster A and C significantly differed with respect to expression levels of 9 genes (Table SIIc). The functional significance of these differences were not subject of further investigation in this study.

We then coupled the profiles to survival data. As shown in Fig. 2a, cluster B gene expression profiles were associated with good prognosis (median survival, defined as time between surgery and death, >6 years; exact value could not be calculated for available follow-up time) whereas gene expression profiles in cluster A and C were associated with median survival of 467 and 135 days, respectively. Results were highly significant between clusters A and B (p<0.0001), A and C (p=0.0078), and C and B (p<0.0001). Survival analysis of the entire cohort, including the 6 non-gliomas, is presented in Fig. SI.

**Fig. 2:**
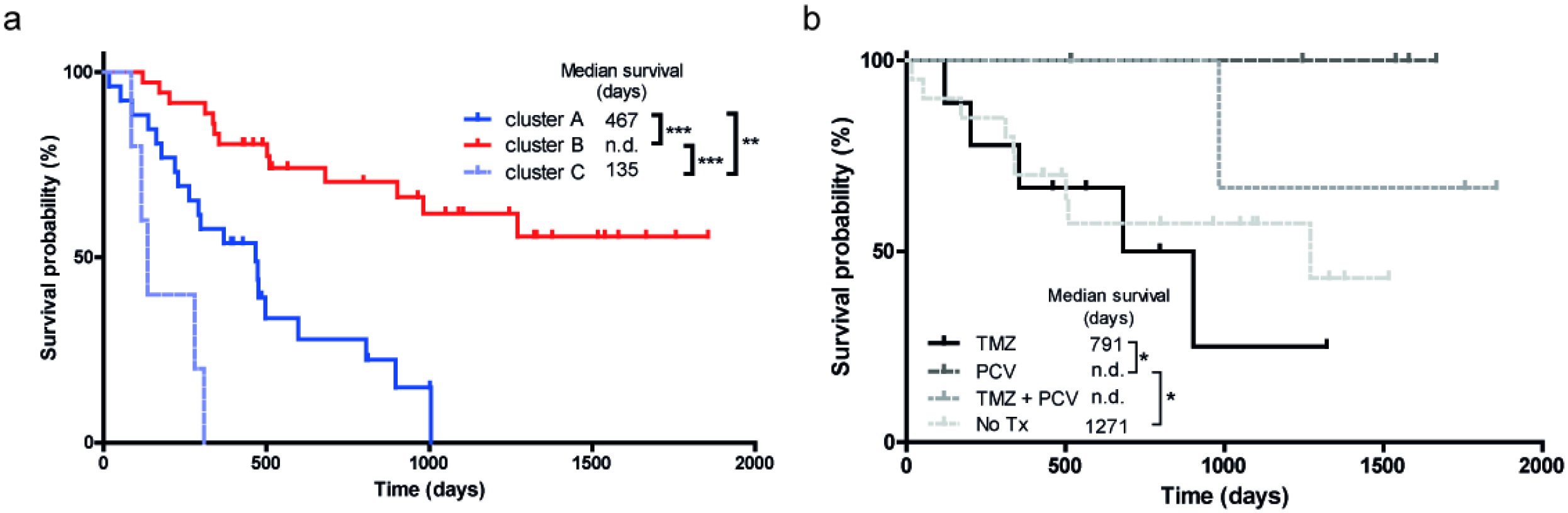
Kaplan-Meier analyses of unsupervised clusters. a) For patients in clusters A (n=26) and C (n=5) median survival was 467 days and 135 days, respectively. For cluster B (n=38) the follow-up period was too short to determine median survival. Survival of patients in cluster B was significantly better than of clusters A and C (both p<0.0001). Survival in cluster A was significantly better than C (P=0.008). b) Oligodendroglioma patients in cluster B (diagnosed according to WHO 2016 classification, and treated with PCV) performed better than astrocytoma patients in this cluster (all treated with TMZ) (p=0.04). Two patients (N16-06 and N16-10) were excluded from analysis because no survival data was present.

### Mutation analysis

We next investigated whether we could identify nucleotide variations that significantly associate with the subgroups. To this end we included all identified sequence variants by base calling with a coverage of >10% of the unique reads. This analysis revealed that a hotspot mutation in the metabolic enzyme isocitrate dehydrogenase 1 (IDH1 p.R132H) mutation [11] was significantly associated with cluster B (p=4E-10 when compared to cluster A, see Table SIII). Also a duplication in succinate dehydrogenase A (SDHA) of unknown significance was strongly highly associated with group B (p=6E-6). The test also identified known oncogenic mutations in mitochondrial IDH2 in four patients (two p.R172K, one p.R172C and one p.R172M mutation, Table SI). Gene expression profiles from these gliomas co-clustered with those of the IDH1 p.R132H gliomas in cluster B (Fig. 1, black in IDH status annotation bar). All IDH1^R132^ and IDH2^R172^ mutations were in retrospect confirmed by routine genetic analyses (not shown). As described before, also other variants of IDH1 (p.V178I, p.Y183C) were identified by the assay [35] (Table SI).

We then analyzed the profiles (generated from treatment-naive tumor samples) in relation to survival with treatment and histopathology as additional parameters. Retrospective analysis of histopathology and clinical follow-up data revealed that all patients in clusters A and C received temozolomide (TMZ) and/or radiotherapy upon signs of tumor progression after surgery. We therefore concentrated on patients in cluster B who were treated with TMZ (n=9, astrocytomas), procarbazine/lomustine/vincristine (PCV) chemotherapy (n=6, all 1p/19q codeleted oligodendrogliomas), both adjuvant therapies (n=3), or did not receive additional treatment (n=19). Of 2 patients adjuvant therapy status was unknown (n=2). As shown in Fig. 2b, outcome for oligodendroglioma patients treated with PCV was better than for astrocytoma patients treated with TMZ (p=0.04).

Whereas group B was dominated by IDH-mutated gliomas, this group also contained 6 IDH wild-type glioblastomas. Comparison of gene expression profiles of these gliomas with those of cluster A revealed that expression levels of all VEGF isoforms were significantly lower in the group B *IDH*^*wt*^ gliomas (Table SIV). This suggests that these 6 tumors grouped with the *IDH*^*mut*^ gliomas based on the lack of an angiogenic response.

### t/RNA-NGS based molecular diagnosis

To investigate whether t/RNA-NGS profiles carry diagnostic information we retrospectively collected routine diagnoses based on histopathology, and investigated correlations. We grouped the gliomas according to the WHO 2016 grading system [41] as grade II/III astrocytomas (n=12), grade II/III oligodendrogliomas (n=19) and glioblastomas (n=38). Based on these groups we performed the Wilcoxon Mann-Whitney U test to identify differentially expressed genes. We identified 79 genes that were differentially expressed between diffuse grade II/III oligodendrogliomas and glioblastomas with high significance (Table SVa). Expression levels of 50 genes were significantly different between grade II/III astrocytomas and glioblastomas (Table SVb), whereas expression levels of only 1 gene (ALK) differed significantly between astrocytomas and oligodendrogliomas (Table SVc). Both oligodendrogliomas and astrocytomas were distinguished from glioblastomas by the IDH1 p.R132H mutation. Among the transcripts that were differentially expressed between grade II/III astrocytomas/oligodendrogliomas and glioblastomas was EGFR^vIII^ that was expressed in 30% of glioblastomas and in none of the grade II/III gliomas (Fig.3a), in good agreement with literature [42]. Expression levels in EGFR^vIII^-positive glioblastomas ranged from 6-1450 (n=13; mean FPM=550) as compared to 0-2.7 in glioblastomas, designated as EGFR negative (n=25; mean FPM=0.16). Expression of wild-type EGFR was found among all gliomas (data not shown), but was more prominent in glioblastomas than in grade II/III gliomas (mean FPM=612 vs. 126; p=0.0061). Levels of EGFR and MET expression in glioblastomas were inversely correlated (Spearman R=−0.59, p<0.0001; Fig. 3b). Not surprising given the strong association of IDH1 mutations and grade II/III gliomas with cluster B of Fig. 1, the supervised analysis again identified ErbB3, ErbB4 and TrkB as potentially targetable proteins in grade II/III gliomas (Fig. 3c).

**Fig. 3:**
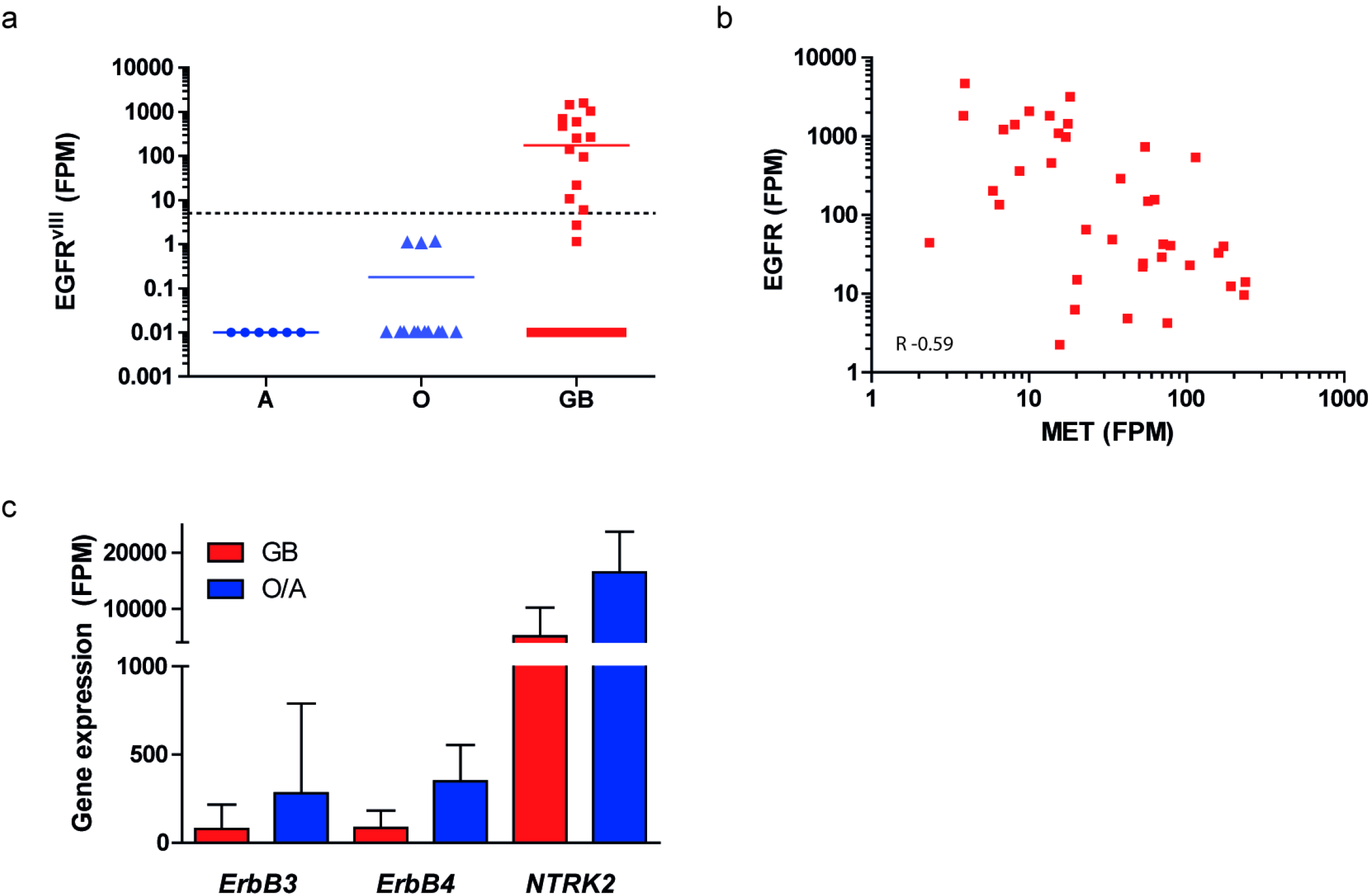
Gene expression in grade II/III astrocytomas, oligodendrogliomas and glioblastoma. a)EGFR^vIII^ was detected at significant levels in 30% of glioblastomas (GB, red) but not in grade II/III astrocytomas (A) and oligodendrogliomas (O) (note the log scale of the Y-axis, expression <5 FPM was considered EGFR^vIII^-negative). b) Wild-type EGFR expression is inversely correlated with expression of MET in glioblastoma (Spearman R= −0.59; p <0.0001). c) Mean expression of *ErbB3, ErbB4,* and *NTRK2* in GB (red) and grade II/III O/A (blue). Error bars indicate the standard deviation.

### Carbonic anhydrase 12 variant 2 is a poor prognostic factor

To investigate whether prognostic factors other than *IDH1*-mutations could be identified, we performed subgroup analysis on tumor profiles from *IDH*^*wt*^ patients with survival <14 months and >14 months. A Fisher’s exact test on the sequence variations in these groups identified a splice isoform of CA12 that lacks exon 9 and expression levels of which is associated with poor survival (CA12v2; Fig. 4a). Variant detection was based on one smMIP with the exon 8-9 and exon 8-10 boundaries in its ROI. CA12v2 was never expressed in *IDH*^*mut*^ gliomas. CA12v2 expression values higher than 50 FPM translated in poor prognosis as shown by Kaplan-Meier analysis (272 vs.1002 days from surgery to death, p=0.0137) (Fig. 4b).

**Fig. 4:**
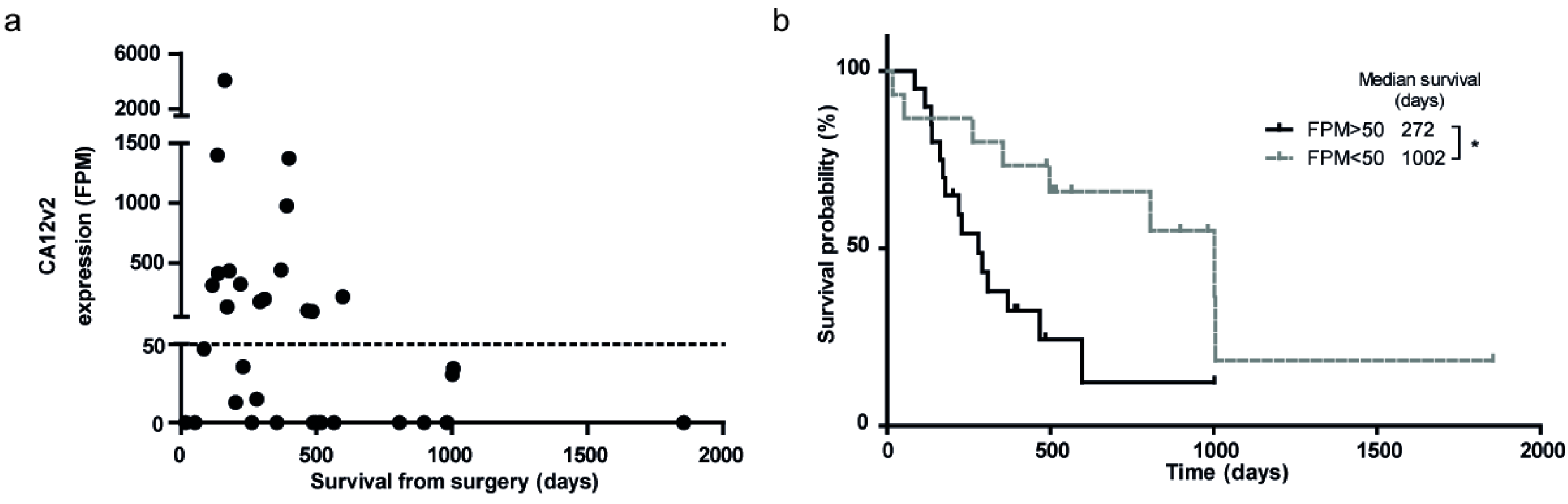
Expression of CA12v2 is a poor prognostic factor. a) CA12v2 expression levels in the positive patient samples included in panel A. Expression >50 FPM is a poor prognostic survival marker. b) Kaplan-Meier curve of IDH^WT^ gliomas, grouped based on CA12v2 expression. Samples with expression levels >50 FPM were considered CA12v2 positive.

### Validation of t/RNA-NGS data

We have previously shown that the quality of t/RNA-NGS data correlates well with w/RNA-NGS [34]. Supervised analysis of gene expression profiles of *IDH1*^*RI32H*^ and *IDH1*^*wt*^ gliomas already revealed a strong correlation of IDH1^R132H^ expression with low expression levels of *BCAT1* and *LDHA* genes, in line with previous reports that demonstrated hypermethylation of the promoters of these genes in *IDH1*-mutated gliomas [35, 43, 44].

The t/RNA-NGS assay was designed to also measure transcripts with involvement in other cancer types. Notably, androgen receptor (AR) was frequently expressed in gliomas at relatively high levels. To confirm and validate our findings further on the proteome level we performed immunohistochemical analyses for EGFR, MET, CA12 and AR on a number of tumors in which t/RNA-NGS data revealed low or high expression levels of the corresponding genes. Transcript levels for EGFR, MET, CA12, and AR correlated with protein levels as established with IHC (Fig. 5). In one case we found that EGFR-transcript levels from frozen tumor tissue did not match with protein levels in FFPE blocks. This discrepancy could be attributed to tumor heterogeneity since the FFPE block contained both EGFR-positive and negative areas (Fig. 5a,b).

**Fig. 5:**
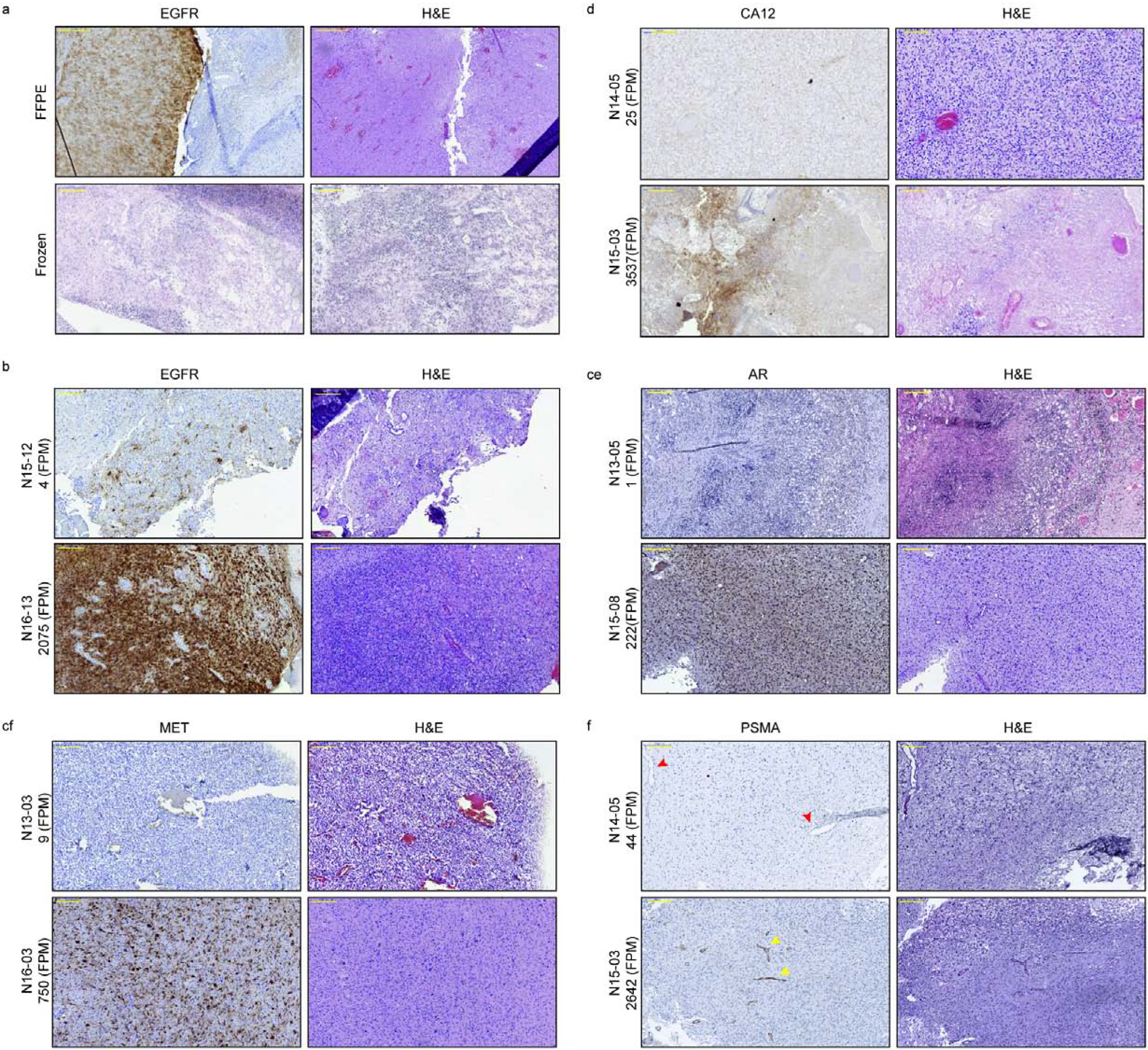
Immunohistochemical staining of EGFR, MET, CA12, AR, and PSMA. a) EGFR staining of tumor N15-12 (negative in the smMIP assay) was negative in the frozen section, but partly positive in the FFPE section, indicating intratumoral heterogeneity of EGFR expression. High EGFR gene expression in tumor N16-13 was consistent with protein expression (b). IHC staining for MET (c), CA12 (d), and AR (e) of tumor samples with low (upper panels) and high (lower panels) expression levels of MET, CA12 and AR respectively in the t/RNA-NGS assay. Expression levels in FPM are stated with the sample name. f) Staining for PSMA showed endothelial staining in gliomas with high PSMA FPM values, as indicated with the yellow arrowheads in the lower panel. In gliomas with low expression (N14-05), no staining was observed (red arrowheads in upper panel). Original magnification for upper panel in A, 5x. For Lower panel of A, and other panels magnification is 10x. Yellow annotated bar for 5x 500μm, for 10x 200μm.

We also detected prominent expression of *FOLH1*, the gene encoding PSMA in approximately 70% of brain tumors of different histological type, including glioblastomas, oligodendrogliomas and astrocytomas (FPM range 0-9148). To validate this finding we performed IHC analysis for PSMA on a number of tumors with either low or high *FOLH1* FPM values. IHC revealed expression of PSMA on the blood vessels of tumors with high *FOLH1* FPM values (Fig. 5f).

### TrkB is expressed as a kinase-defective protein in IDH^mut^ gliomas

An intriguing finding was that *NTRK2*, the gene encoding TrkB, was expressed at relatively high levels in IDH1 p.R132H mutated gliomas (Fig. 3). Upon closer examination, we found a dysbalanced reaction of the 5 smMIPs that were designed against different ROIs of the *NTRK2* transcript, with the smMIPs designed to sequence ROIs in the 3’-end yielding lower number of reads than smMIPs on ROIs in the 5’-end of the transcript (Fig. 6a). To investigate whether this is due to low efficacy of the corresponding smMIPs, or whether this is related to dysbalanced expression of the corresponding exons, we chose a set of 8 *IDH1*^*R132H*^ mutated gliomas with high *NTRK2* expression levels (FPM 23885-28785) and 8 *IDH1*^*wt*^ brain tumors with low to intermediate *NTRK2* expression levels (FPM 30-3011) and performed whole transcriptome RNA-NGS. Results confirmed the t/RNA-NGS expression levels for *NTRK2* in all samples (Fig. 6b). Annotation of the data confirmed the predominant expression of a truncated transcript lacking exons 15-21 (containing the kinase domain) in all cancers analyzed (Fig. 6c). The link between the IDH1 p.R132H and this specific NTRK2 variant is unclear.

**Fig. 6:**
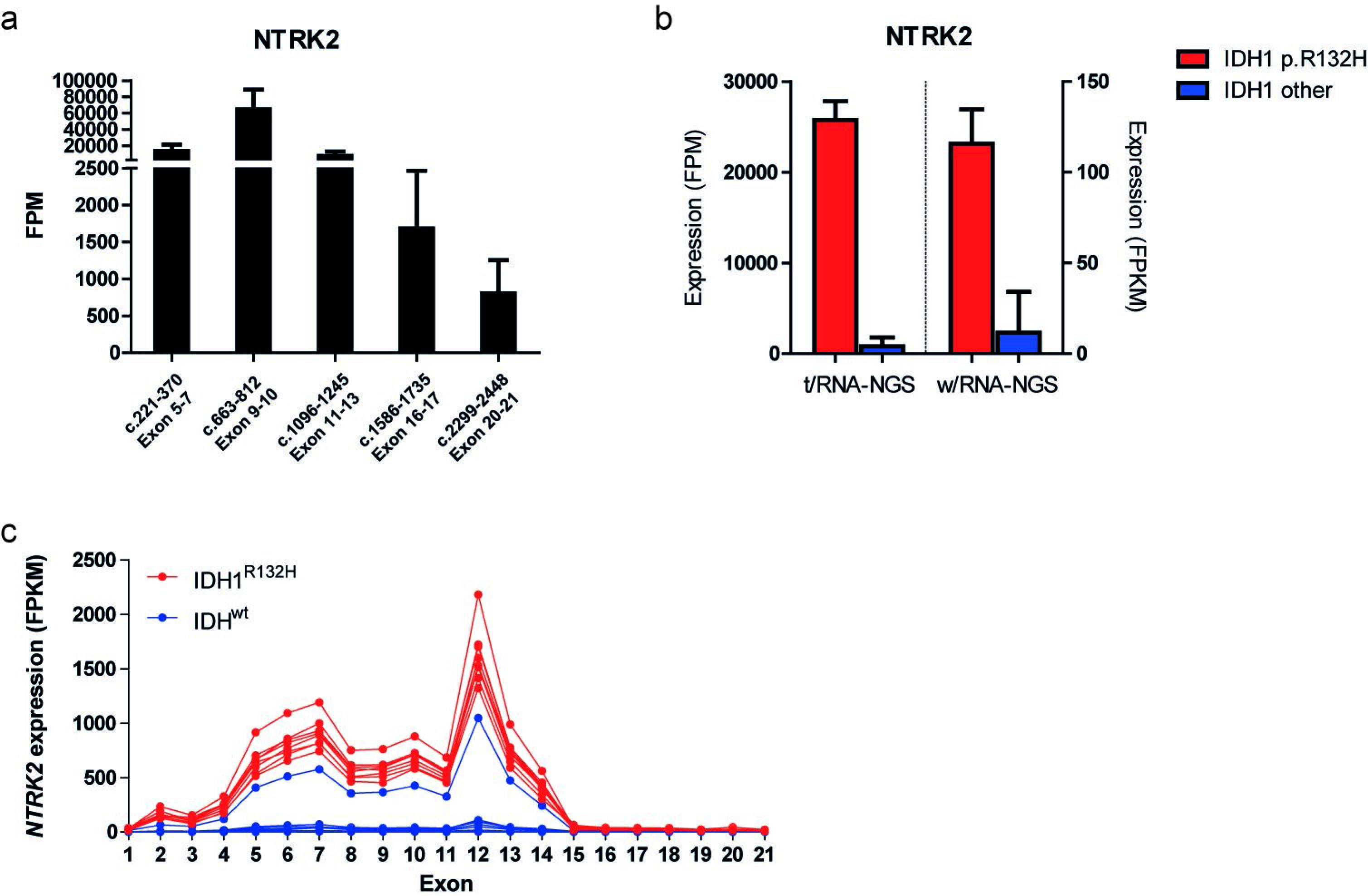
NTRK2 expression in brain tumors. a) FPM expression levels of the 5 smMIPs targeting NTRK2 in IDH1 p.R132H gliomas (n=28, mean±SD). smMIPs at the 5’ end of the transcript show higher FPM levels than smMIPs at the 3’ end of the transcript. b) Mean expression of NTRK2 in 8 IDH1 p.R132H-mutated gliomas, and 8 brain tumors without the IDH1 p.R132H mutation (IDH1 other). On the left y-axis t/RNA-NGS FPM levels are shown, while on the right y-axis w/RNA-NGS FPKM levels are depicted (mean±SD). c) Exon-specific w/RNA-NGS FPKM expression levels for *NTRK2*, showing absence of exons 15-21 in all NTRK2-positive gliomas.

### Actionable markers in individual brain cancers

In the above group-based analyses we concentrated on grade II-IV gliomas. From profiles of individual cancers also interesting observations were made. In the ganglioglioma a *BRAF*^*V600E*^ mutation was found which was confirmed via genetic testing (Table S1). This mutation has been described before in this tumor type [45]. Gangliogliomas are mostly slow growing, low-grade gliomas but some patients will undergo malignant progression. Mutations in BRAF have been identified as poor prognostic factors for ganglioglioma [46].

Other observations were that the brain metastasis of lung carcinoma expressed high levels of VEGFR2 and VEGF-isoforms, suggesting an angiogenic phenotype. Also, MET was expressed at relatively high levels whereas EGFR expression was low. The ependymoma showed high levels of EGFR^vIII^ In the DNET, considered as a benign grade I tumor, we found high expression levels of ErbB3, FGFR2, FOLH1 and AXL.

### Analysis of recurrent tumors

Our cohort included a primary and recurrent tumor from one patient. This patient underwent surgery 4 months after first clinical presentation of an *IDH* mutated grade II diffuse astrocytoma. A wait-and-see policy was applied and the patient received a second debulking operation after signs of progression 2.5 years after first surgery. The recurrent tumor was diagnosed as an *IDH* mutated grade III anaplastic astrocytoma. Correlation of the profiles from primary and recurrent tumors was highly significant (Fig. 7, Spearman R=0.96; p<0.0001). AR was downregulated in the recurrent tumor compared to the primary tumor, while CBR1, G6PC and FOLH1 were upregulated.

**Fig. 7:**
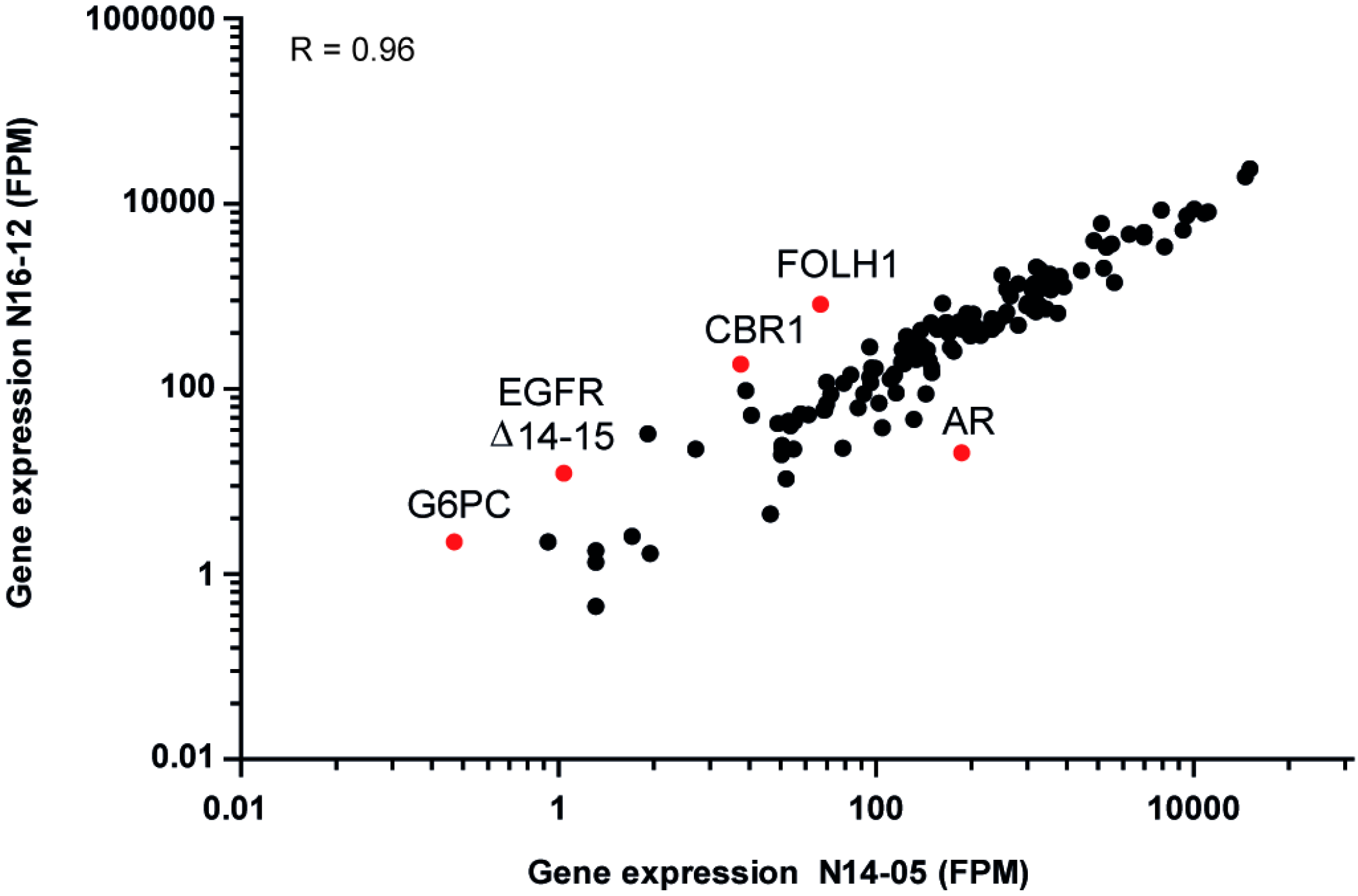
Correlation of gene expression of a primary and recurrent tumor of one patient. t/RNA-NGS profiles are highly correlated (Spearman R=0.96, p<0.0001). Genes of which expression changed 10-fold between both profiles were considered significant (red dots). CBR1, FOLH1, and G6PC were upregulated while AR was downregulated in the recurrent tumor. X-axis: primary tumor, Y axis: recurrence.

## Discussion

We here describe the clinical application of a relatively new and highly cost-effective hybrid multiplex Q-PCR-next generation sequencing assay. The assay generates expression profiles and sequence information of genes that have been identified by literature surveys as potentially important for diagnosis, prognosis, but also for potential prediction of response to precision medicines in a variety of cancer types. New smMIPs for the detection of newly identified transcripts or mutations can be added to the existing panel at low cost.

We here tested the technique on a cohort of brain cancers, of which 69 gliomas, a tumor type characterized by high molecular heterogeneity and in desperate need for alternative treatment. Much has been learned in the last decades on the molecular drivers of gliomagenesis and contributors to tumor progression, a result of high throughput genomic and transcriptomic sequence analyses that have been collected in open access databases such as TCGA. Whereas these databases pinpoint molecular drivers in gliomas as a group, the biology in individual cancers is mostly a black box. In the routine clinical setting in most hospitals molecular diagnosis of glioma is implemented as add-on to histopathology to aid diagnosis and prognosis, but influence on therapeutic decision making is limited. Most important biomarkers include IDH mutations, conferring relatively good prognosis, and loss of chromosome arms 1p and 19q, leading to a diagnosis of oligodendroglioma if combined with an *IDH* mutation (predicting a better response to PCV treatment) and methylation of the *MGMT* promoter (predicting better response to TMZ) [47–49]. *EGFR* status is also determined, amplification of which, in combination with the EGFR^vIII^ mutation, is associated with glioblastoma [50]. Because genes do not necessarily need to be mutated to endow them with hyperactivity, we anticipated that molecular profiling should concentrate on RNA rather than DNA.

Diffuse gliomas of the brain present with different molecular mechanisms involved in gliomagenesis. Whereas the majority of gliomas of grade II and III start with mutations in *IDH1* or *IDH2*, high grade gliomas present with genetic aberrations on different levels of cell signaling, including aberrant expression of membrane tyrosine kinases, mutations in downstream signaling molecules PIK3CA and inactivating mutations in the phosphatase PTEN and *CDKN2A* [11, 51]. Often, multiple aberrations co-occur in individual tumors, whereas precision medicines often use only one biomarker. For instance, MET overexpression may lead to inclusion of patients for treatment with MET-inhibitors, whereas such treatment can lead to rapid selection of subclones that overexpress rescue kinases, or overexpress inhibitors of phosphatases [52–54]. It is therefore of high importance that methods become available that give a comprehensive overview of aberrant molecular pathways in individual cancers that can aid in patient stratification for precision medicines. We here show that t/RNA-NGS gives important insights in the mechanisms operating in individual gliomas. The strong patterns in the gene expression data and its correlation with mutation status and survival suggests that tumor-specific gene expression signatures in individual gliomas are relevant to designing therapy for this disease.

Our data confirm the high intertumoral heterogeneity in gliomas and reveal tumor targets in subsets of gliomas that are associated with other cancer types, opening new avenues for targeted therapy. Our data confirm expression of the prostate cancer marker PSMA on microvasculature of glioblastoma, possibly providing opportunities for tumor-vascular targeting [55–57]. In a recent publication high expression levels of androgen receptor were described in glioblastomas, that actually translated into sensitivity to the AR antagonist enzalutamide [39]. Notably, our data also confirmed the identification of CA12v2 as a marker of poor prognosis in glioblastoma patients [58]. CA12 is a transmembrane enzyme that is involved in intracellular pH homeostasis and extracellular acidification, providing a biological explanation for an association with poor survival in glioma and other cancers [59]. How the lack of exon 9 in CA12v2 affects its function is not known. Because CA12v2 is not expressed in normal brain, it may be a good target for glioblastomas expressing this variant [60]. Other interesting therapeutic targets for low-grade gliomas that were identified by our assay are NTRK2, ERBB3 and ERBB4.

In conclusions we here present a novel t/RNA-NGS ‘one size fits all’ assay that can provide a molecular diagnosis and prognosis for gliomas and possibly other cancers in an unsupervised analysis. The advantages of this technique as compared to conventional whole RNA-seq are multiple: it generates quantitative gene expression values at low costs and quantitatively detects alternative splicing by direct sequencing of exon-exon boundaries. Because of the targeted nature of the assay, Illumina sequencing gives higher coverage per ROI than whole RNA-seq providing more reliable mutation analysis. In our cohort of brain tumors, IDH and BRAF mutations were found that were all confirmed using routine DNA sequencing. The assay gives insight in metabolic pathways [35] and tyrosine kinase signaling pathways and the probeset can be expanded with new smMIPs for sequencing of other ROIs. A disadvantage is the targeted nature of the assay, requiring in depth preliminary knowledge of expected aberrations. An issue that cannot be controlled on forehand at this moment is the efficacy of individual smMIPs. If an ROI with a mutation hotspot is not efficiently covered, such mutations will be missed. This risk can however be assessed by monitoring the absolute read numbers of the corresponding smMIPs.

Because the assay detects expression of genes, involved in multiple pathways that have involvement in multiple cancer types, t/RNA-NGS may allow repurposing of drugs in an individualized manner.

## Materials and Methods

The aim of this study was to identify the value of targeted RNA-NGS for establishing a molecular diagnosis and prognosis for individual brain cancer patients, and to identify novel biomarkers for glioma. To this end we performed targeted RNA-NGS profiling of 75 brain tumors in a retrospective setting and analyzed the results in connection with histopathological and genetic routine diagnosis and clinical outcome.

### Patients

All studies described here were performed with brain tumor tissue from patients who were operated upon between 2013 and 2017 (n=75). Researchers were blinded to histopathology and clinical outcome. The study protocol was approved by the Ethical Committee for Human Experimentation of the Radboudumc. All patients signed informed consent. Directly after surgery, tissue samples were snap frozen in liquid nitrogen and stored at −80°C until further processing. In retrospect, patient characteristics were extracted from Radboudumc electronic patient files (EPIC) and documented in the electronic data capture system CASTOR. Histopathology and molecular diagnoses were extracted from the Dutch Pathology archive PALGA.

### *RNA preparation* and cDNA synthesis

Cryosections of 10 μm were cut for RNA isolation using TRIzol (ThermoFisher Scientific, Waltham, MA). For every sample a 4 μm serial section was stained with H&E to estimate percentage tumor area by an experienced neuropathologist (BK). RNA was reverse transcribed into cDNA using random hexamer primers and Superscript II (Invitrogen, CA) according to standard protocols.

In parallel, tissue was processed to formalin-fixed paraffin-embedded (FFPE) tissue blocks that were used for routine diagnosis. Samples from 2014 onwards were also subjected to genetic analysis using targeted DNA next generation sequencing [18].

### T/RNA-NGS with smMIPs

All enzymes were from NEB (Ipswich, MA) unless stated otherwise. The procedure of smMIP-based targeted RNA Next Generation Sequencing (t/RNA-NGS) to detect expression of metabolic genes has been described before [34–36]. SmMIPs were designed by an adjusted version of the MIPgen algorithm [37] and were ordered from Integrated DNA Technologies (Leuven, Belgium). The set of smMIP probes used in [34] was expanded with probes for detection of novel transcripts of interest that play a role in a wide variety of cancer types, including prostate cancer markers prostate specific membrane antigen (PSMA [38]) and androgen receptor and its splice variant ARv7 [39](Table 1). Furthermore, smMIP probes were included for specific detection of splice variants by placing extension and ligation probes on neighboring exons, e.g. probes on exons 1/8 of Epithelial Growth Factor Receptor (EGFR) for detection of EGFR^vIII^ [36]. Probes for detection of Vascular Endothelial Growth Factor (VEGF) isoforms 121, 165 and 189 were designed with extension and ligation probes in exons 5/8, 5/7 and 5/6, respectively. Additional smMIPs were included that were directed against shared exons of transcript variants. For transcripts in which relevant mutations can be expected, smMIPs were selected with extension and ligation probes flanking the respective region of interest (ROI). Based on intrinsic variations in smMIP efficacy, we included at least 5 smMIPs for each transcript of interest, evenly distributed along the transcript. cDNA was subjected to smMIP capture as described in [34]. In short, a mixture of 936 different smMIPs (phosphorylated using T4-polynucleotide kinase) was hybridized to 50 ng of cDNA for each patient sample. Each smMIP molecule is designed to hybridize via its extension and ligation probes in an inverted manner to its target cDNA ROI, leaving a gap of maximum 112 bases between the extension and ligation probe [37]. After overnight enzymatic gap-filling (KlenTaq polymerase, Epicentre, Madison, WI) and ligation (Ampligase, Epicentre) circular smMIPs are formed, the number of which is linearly related to the number of RNA molecules present in the original sample. After treatment with exonuclease to remove cDNA and linear, non-reacted smMIPs the libraries of circular smMIPs were subjected to PCR with barcoded primers. After purification of the correct 266 bp PCR products with Ampure beads (Beckmann Coulter Genomics, High Wycombe, UK) and verification of correct PCR product size of 266 bp on a TapeStation 2200 (Agilent Technologies, Santa Clara, CA), PCR products were sequenced on the Illumina Nextseq platform (Illumina, San Diego, CA) at the Radboudumc sequencing facility (output 2×151 bases). The barcode in the PCR primers allows for pooling of multiple samples into one sequencing library and de-multiplexing of reads after sequencing.

### Data processing

For each patient sample, all reads were mapped against reference transcripts (UCSC human genome assembly hg19) using the SeqNext module of JSI SequencePilot version 4.2.2 build 502 (JSI Medical Systems, Ettenheim, Germany). Since all individual smMIP probes contain a unique random octamer sequence adjacent to the ligation probe (unique molecular identifier or UMI), all identical PCR products can be assigned to one originating circularized (unique) smMIP, excluding PCR amplification bias and making the assay quantitative. The number of unique reads for each individual smMIP probe was divided by the total number of unique reads within the sample and multiplied by a million, resulting in a fragment per million (FPM) value for each smMIP. Mean FPM values from all smMIPs covering different ROIs in the same transcript were considered to represent gene expression levels in the tumor sample. Using the base calling algorithm of SeqNext a list of single nucleotide variations and insertions and deletions was generated for each sample.

### Data analysis and statistics

Unsupervised hierarchical cluster analysis with the gene expression data was performed using R (version 3.4.3) programming, using log-transformed gene expression values (mean FPM + 0.01 to prevent a log 0 transformation), Manhattan distance to calculate the distance between gene expression profiles, and the group average method for agglomerative clustering (Unweighted Pair Group Method with Arithmetic Mean). The Wilcoxon Mann-Whitney U test was used to find differentially expressed genes between clusters. Associations of mutations with clusters were calculated using Fisher’s exact test. Multiple testing corrections were done using Benjamini Hochberg (FDR<0.05).

Kaplan-Meier survival curves of 75 brain cancer patients were generated using Graphpad Prism, version 5.03. Patients who were still alive at the date of analysis or were lost to follow-up, were censored in the survival data. P-values were calculated using the Log-Rank test. All p-values are indicated as *(p<0.05), ** (p<0.01), *** (p<0.001), **** (p<0.0001), unless specified otherwise.

### Immunohistochemistry

Immunohistochemistry (IHC) was performed on 4 μm sections of FFPE tissue or on frozen sections, adjacent to those used for t/RNA-NGS. Frozen sections were air dried and fixed with 4% paraformaldehyde (PFA) solution for 20 min at room temperature before staining. After appropriate epitope retrieval (for FFPE sections), antibodies rabbit-anti-Prostate-Specific Membrane Antigen (PSMA) (Abcam; ab133579), rabbit-anti-Carbonic Anhydrase 12 (CA12) (Sigma Life Sciences; HPA008773), mouse-anti-androgen receptor (AR) (Santa Cruz; sc-7305), rabbit-anti-MET (Cell Signaling Technologies; #8198), and rabbit-anti-EGFR (Cell Signaling Technologies; #4267) were used. Sections were incubated with primary antibody in normal antibody diluent (Immunologic, Duiven, The Netherlands) overnight at 4°C. Primary antibody detection was done using BrightVision polyHRP-anti-rabbit IgG (Immunologic, Duiven, The Netherlands), or BrightVision polyHRP-anti-mouse/rabbit/rat IgG (Immunologic, Duiven, the Netherlands) for AR staining. Sections were counterstained with haematoxylin and mounted with Quick-D mounting medium (Klinipath, Duiven, The Netherlands). As control staining, secondary antibody-only stainings were performed.

### Whole transcriptome RNA-NGS (w/RNA-NGS)

Total RNA was isolated using the Qiagen AllPrep DNA/RNA Mini Kit following the manufacturer’s protocol for animal tissues (Qiagen, Hilden, Germany). RNA sequencing libraries were prepared from gliomas with high and low *NTRK2* expression levels (n=8 for each group) using the KAPA RNA HyperPrep Kit with RiboErase (HMR) (KAPA Biosystems, Wilmington, MA) following the manufacturer’s protocol. Briefly, 100ng of total RNA was used to generate libraries, which were fragmented for 8 minutes at 94°C. Adapter stock concentration used was 750nM and libraries were amplified for 11 cycles. Duplex “Y” adapter sequences with molecular barcodes were generated by IDT (Integrated DNA Technologies, Skokie, Illinois). Final libraries were quantified on a High Sensitivity Bioanalyzer chip (Agilent, Santa Clara, CA) and sequenced at 1.4pM with 10% PhiX on the Illumina NextSeq 550 Sequencer (Illumina, San Diego, CA). Raw FASTQ files were mapped to the human genome (hg19) with STAR (v2.5.3) aligner [40]. Mapped reads were filtered and deduplicated using sambamba v0.6.6 and feature quantification was performed using featureCounts v1.5.0-p1 against the RefSeq database (downloaded from the UCSC genome browser on 01/05/2017). Gene and exon-level fragments per kilobase per million mapped reads (FPKMs) were calculated using a custom python script.

**Table 1: Metabolic transcripts for smMIP design.** SmMIPs were designed against the antisense strand of predicted transcritps (UCSC human genome assembly hg19 and splice variant specific FASTA sequences). The genes listed are newly added to a previously published panel [34].

## Acknowledgements

We thank Carlijn van de Water and Tessa de Bitter (Radboudumc, Dept. of Pathology), for help with smMIP design. We thank Sanne van Lith (Radboudumc, Dept. of Radiology and nuclear medicine) and Kiek Verrijp (Radboudumc, Dept. of Pathology) for immunohistochemical stainings. KL was funded by the Dutch Cancer Society, grant UvA2014-6839, CvdH was supported by Eurostars (E9616) and the Stop Brain Tumor Foundation.

## References

1. Baugh EH, Ke H, Levine AJ, Bonneau RA, Chan CS: Why are there hotspot mutations in the TP53 gene in human cancers? Cell Death Differ. 2018; 25(1):154–60.

2. Basu S, Murphy ME: Genetic Modifiers of the p53 Pathway. Cold Spring Harb Perspect Med. 2016; 6(4):a026302.

3. Bhattacharya P, Patel TN: Microsatellite Instability and Promoter Hypermethylation of DNA repair genes in Hematologic Malignancies: a forthcoming direction toward diagnostics. Hematology. 2018; 23(2):77–82.

4. Du Z, Lovly CM: Mechanisms of receptor tyrosine kinase activation in cancer. Mol Cancer. 2018; 17(1):58.

5. Hendriks W, Bourgonje A, Leenders W, Pulido R: Proteinaceous Regulators and Inhibitors of Protein Tyrosine Phosphatases. Molecules. 2018; 23(2).

6. Choudhry H, Harris AL: Advances in Hypoxia-Inducible Factor Biology. Cell Metab. 2018; 27(2):281–98.

7. Petrova V, Annicchiarico-Petruzzelli M, Melino G, Amelio I: The hypoxic tumour microenvironment. Oncogenesis. 2018; 7(1):10.

8. Potente M, Gerhardt H, Carmeliet P: Basic and therapeutic aspects of angiogenesis. Cell. 2011; 146(6):873–87.

9. Ariazi EA, Jordan VC: Estrogen-related receptors as emerging targets in cancer and metabolic disorders. Curr Top Med Chem. 2006; 6(3):203–15.

10. Gaillard-Moguilewsky M: Pharmacology of antiandrogens and value of combining androgen suppression with antiandrogen therapy. Urology. 1991; 37(2 Suppl):5–12.

11. Lenting K, Verhaak R, Ter Laan M, Wesseling P, Leenders W: Glioma: experimental models and reality. Acta Neuropathol. 2017; 133(2):263–82.

12. Pant S, Hubbard J, Martinelli E, Bekaii-Saab T: Clinical update on K-Ras targeted therapy in gastrointestinal cancers. Crit Rev Oncol Hematol. 2018; 130:78–91.

13. Agianian B, Gavathiotis E: Current Insights of BRAF Inhibitors in Cancer. J Med Chem. 2018; 61(14):5775–93.

14. Rubin MA, Demichelis F: The Genomics of Prostate Cancer: emerging understanding with technologic advances. Mod Pathol. 2018; 31(S1):S1–11.

15. Mondesir J, Willekens C, Touat M, de Botton S: IDH1 and IDH2 mutations as novel therapeutic targets: current perspectives. Journal of blood medicine. 2016; 7:171–80.

16. Nicolaides TP, Li H, Solomon DA, Hariono S, Hashizume R, Barkovich K et al: Targeted therapy for BRAFV600E malignant astrocytoma. Clin Cancer Res. 2011; 17(24):7595–604.

17. Hyman DM, Piha-Paul SA, Won H, Rodon J, Saura C, Shapiro GI et al: HER kinase inhibition in patients with HER2-and HER3-mutant cancers. Nature. 2018; 554(7691):189–94.

18. Eijkelenboom A, Kamping EJ, Kastner-van Raaij AW, Hendriks-Cornelissen SJ, Neveling K, Kuiper RP et al: Reliable Next-Generation Sequencing of Formalin-Fixed, Paraffin-Embedded Tissue Using Single Molecule Tags. J Mol Diagn. 2016; 18(6):851–63.

19. Neveling K, Mensenkamp AR, Derks R, Kwint M, Ouchene H, Steehouwer M et al: BRCA Testing by Single-Molecule Molecular Inversion Probes. Clin Chem. 2017; 63(2):503–12.

20. Gorgannezhad L, Umer M, Islam MN, Nguyen NT, Shiddiky MJA: Circulating tumor DNA and liquid biopsy: opportunities, challenges, and recent advances in detection technologies. Lab Chip. 2018; 18(8):1174–96.

21. Capper D, Jones DTW, Sill M, Hovestadt V, Schrimpf D, Sturm D et al: DNA methylation-based classification of central nervous system tumours. Nature. 2018; 555(7697):469–74.

22. Brien GL, Valerio DG, Armstrong SA: Exploiting the Epigenome to Control Cancer-Promoting Gene-Expression Programs. Cancer Cell. 2016; 29(4):464–76.

23. Capper D, Stichel D, Sahm F, Jones DTW, Schrimpf D, Sill M et al: Practical implementation of DNA methylation and copy-number-based CNS tumor diagnostics: the Heidelberg experience. Acta Neuropathol. 2018; 136(2):181–210.

24. Kangsamaksin T, Tattersall IW, Kitajewski J: Notch functions in developmental and tumour angiogenesis by diverse mechanisms. Biochem Soc Trans. 2014; 42(6):1563–8.

25. Saharinen P, Eklund L, Pulkki K, Bono P, Alitalo K: VEGF and angiopoietin signaling in tumor angiogenesis and metastasis. Trends Mol Med. 2011; 17(7):347–62.

26. Kusters B, de Waal RM, Wesseling P, Verrijp K, Maass C, Heerschap A et al: Differential effects of vascular endothelial growth factor A isoforms in a mouse brain metastasis model of human melanoma. Cancer Res. 2003; 63(17):5408–13.

27. Frampton GM, Ali SM, Rosenzweig M, Chmielecki J, Lu X, Bauer TM et al: Activation of MET via diverse exon 14 splicing alterations occurs in multiple tumor types and confers clinical sensitivity to MET inhibitors. Cancer Discov. 2015; 5(8):850–9.

28. Lowenstein PR, Castro MG: The value of EGFRvIII as the target for glioma vaccines. Am Soc Clin Oncol Educ Book. 2014:42–50.

29. Navis AC, van Lith SA, van Duijnhoven SM, de Pooter M, Yetkin-Arik B, Wesseling P et al: Identification of a novel MET mutation in high-grade glioma resulting in an auto-active intracellular protein. Acta Neuropathol. 2015; 130(1):131–44.

30. Greenall SA, Johns TG: EGFRvIII: the promiscuous mutation. Cell Death Discov. 2016; 2:16049.

31. Sottoriva A, Spiteri I, Piccirillo SG, Touloumis A, Collins VP, Marioni JC et al: Intratumor heterogeneity in human glioblastoma reflects cancer evolutionary dynamics. Proc Natl Acad Sci U S A. 2013; 110(10):4009–14.

32. Stupp R, Mason WP, van den Bent MJ, Weller M, Fisher B, Taphoorn MJ et al: Radiotherapy plus concomitant and adjuvant temozolomide for glioblastoma. N Engl J Med. 2005; 352(10):987–96.

33. Claes A, Idema AJ, Wesseling P: Diffuse glioma growth: a guerilla war. Acta Neuropathol. 2007; 114(5):443–58.

34. de Bitter T, van de Water C, van den Heuvel C, Zeelen C, Eijkelenboom A, Tops B et al: Profiling of the metabolic transcriptome via single molecule molecular inversion probes. Sci Rep. 2017; 7(1):11402.

35. Lenting K, Khurshed M, Peeters TH, van den Heuvel C, van Lith SAM, de Bitter T et al: Isocitrate dehydrogenase 1-mutated human gliomas depend on lactate and glutamate to alleviate metabolic stress. FASEB J. 2018:fj201800907RR.

36. van den Heuvel C, Das AI, de Bitter T, Simmer F, Wurdinger T, Molina-Vila MA et al: Quantification and localization of oncogenic receptor tyrosine kinase variant transcripts using molecular inversion probes. Sci Rep. 2018; 8(1):7072.

37. O’Roak BJ, Vives L, Fu W, Egertson JD, Stanaway IB, Phelps IG et al: Multiplex targeted sequencing identifies recurrently mutated genes in autism spectrum disorders. Science. 2012; 338(6114):1619–22.

38. Kunikowska J, Bartosz K, Leszek K: Glioblastoma multiforme: another potential application for (68)Ga-PSMA PET/CT as a guide for targeted therapy. European journal of nuclear medicine and molecular imaging. 2018; 45(5):886–87.

39. Zalcman N, Canello T, Ovadia H, Charbit H, Zelikovitch B, Mordechai A et al: Androgen receptor: a potential therapeutic target for glioblastoma. Oncotarget. 2018; 9(28):19980–93.

40. Dobin A, Davis CA, Schlesinger F, Drenkow J, Zaleski C, Jha S et al: STAR: ultrafast universal RNA-seq aligner. Bioinformatics. 2013; 29(1):15–21.

41. Louis DN, Perry A, Reifenberger G, von Deimling A, Figarella-Branger D, Cavenee WK et al: The 2016 World Health Organization Classification of Tumors of the Central Nervous System: a summary. Acta Neuropathol. 2016; 131(6):803–20.

42. Gan HK, Kaye AH, Luwor RB: The EGFRvIII variant in glioblastoma multiforme. J Clin Neurosci. 2009; 16(6):748–54.

43. Tonjes M, Barbus S, Park YJ, Wang W, Schlotter M, Lindroth AM et al: BCAT1 promotes cell proliferation through amino acid catabolism in gliomas carrying wild-type IDH1. Nat Med. 2013; 19(7):901–08.

44. Chesnelong C, Chaumeil MM, Blough MD, Al-Najjar M, Stechishin OD, Chan JA et al: Lactate dehydrogenase A silencing in IDH mutant gliomas. Neuro Oncol. 2014; 16(5):686–95.

45. Stone TJ, Keeley A, Virasami A, Harkness W, Tisdall M, Izquierdo Delgado E et al: Comprehensive molecular characterisation of epilepsy-associated glioneuronal tumours. Acta neuropathologica. 2018; 135(1):115–29.

46. Dahiya S, Haydon DH, Alvarado D, Gurnett CA, Gutmann DH, Leonard JR: BRAF(V600E) mutation is a negative prognosticator in pediatric ganglioglioma. Acta neuropathologica. 2013; 125(6):901–10.

47. Chen Y, Hu F, Zhou Y, Chen W, Shao H, Zhang Y: MGMT promoter methylation and glioblastoma prognosis: a systematic review and meta-analysis. Archives of medical research. 2013; 44(4):281–90.

48. Hegi ME, Stupp R: Withholding temozolomide in glioblastoma patients with unmethylated MGMT promoter–still a dilemma? Neuro Oncol. 2015; 17(11):1425–7.

49. Molenaar RJ, Verbaan D, Lamba S, Zanon C, Jeuken JW, Boots-Sprenger SH et al: The combination of IDH1 mutations and MGMT methylation status predicts survival in glioblastoma better than either IDH1 or MGMT alone. Neuro Oncol. 2014; 16(9):1263–73.

50. Phillips AC, Boghaert ER, Vaidya KS, Mitten MJ, Norvell S, Falls HD et al: ABT-414, an Antibody-Drug Conjugate Targeting a Tumor-Selective EGFR Epitope. Mol Cancer Ther. 2016; 15(4):661–9.

51. Sibin MK, Bhat DI, Narasingarao KV, Lavanya C, Chetan GK: CDKN2A (p16) mRNA decreased expression is a marker of poor prognosis in malignant high-grade glioma. Tumour Biol. 2015; 36(10):7607–14.

52. Birkman EM, Elzagheid A, Jokilehto T, Avoranta T, Korkeila E, Kulmala J et al: Protein phosphatase 2A (PP2A) inhibitor CIP2A indicates resistance to radiotherapy in rectal cancer. Cancer Med. 2018; 7(3):698–706.

53. Navis AC, Bourgonje A, Wesseling P, Wright A, Hendriks W, Verrijp K et al: Effects of dual targeting of tumor cells and stroma in human glioblastoma xenografts with a tyrosine kinase inhibitor against c-MET and VEGFR2. PLoS One. 2013; 8(3):e58262.

54. van den Heuvel C, Navis AC, de Bitter T, Amiri H, Verrijp K, Heerschap A et al: Selective MET Kinase Inhibition in MET-Dependent Glioma Models Alters Gene Expression and Induces Tumor Plasticity. Molecular cancer research: MCR. 2017; 15(11):1587–97.

55. Nomura N, Pastorino S, Jiang P, Lambert G, Crawford JR, Gymnopoulos M et al: Prostate specific membrane antigen (PSMA) expression in primary gliomas and breast cancer brain metastases. Cancer Cell Int. 2014; 14(1):26.

56. Wernicke AG, Edgar MA, Lavi E, Liu H, Salerno P, Bander NH et al: Prostate-specific membrane antigen as a potential novel vascular target for treatment of glioblastoma multiforme. Arch Pathol Lab Med. 2011; 135(11):1486–9.

57. Salas Fragomeni RA, Menke JR, Holdhoff M, Ferrigno C, Laterra JJ, Solnes LB et al: Prostate-Specific Membrane Antigen-Targeted Imaging With [18F]DCFPyL in High-Grade Gliomas. Clin Nucl Med. 2017; 42(10):e433–e35.

58. Haapasalo J, Hilvo M, Nordfors K, Haapasalo H, Parkkila S, Hyrskyluoto A et al: Identification of an alternatively spliced isoform of carbonic anhydrase XII in diffusely infiltrating astrocytic gliomas. Neuro-oncology. 2008; 10(2):131–8.

59. Mboge MY, Mahon BP, McKenna R, Frost SC: Carbonic Anhydrases: Role in pH Control and Cancer. Metabolites. 2018; 8(1).

60. Fiedler L, Kellner M, Gosewisch A, Oos R, Boning G, Lindner S et al: Evaluation of (177)Lu-CHX-A”-DTPA-6A10 Fab as a radioimmunotherapy agent targeting carbonic anhydrase XII. Nucl Med Biol. 2018; 60:55–62.

